# Acoustic Recognition of Individual Animals in the Presence of Unknown Individuals

**DOI:** 10.1101/2024.12.18.629284

**Authors:** Lifi Huang, Rohan Clarke, Daniella Teixeira, André Chiaradia, Bernd Meyer

## Abstract

Passive acoustic monitoring is firmly established as an effective non-invasive technique for wildlife monitoring. The analysis of animal vocalizations recorded in their natural habitats is commonly used to monitor species occupancy, distribution mapping and community composition. However, the ability to distinguish between individual animals by their vocalisations remains under-explored and presents an exciting opportunity to study individual animal behavior and population demographics in more detail. In this work, we investigate bioacoustic individual-level recognition. In contrast to existing work, we focus on settings where only a subset of the existing population is known and labeled. This is crucial because wildlife populations are constantly changing so that solutions operating only within a known set of individuals are not realistically applicable in the wild. Using two novel datasets, we show that models initially trained to classify only known individuals can also be extended to detect new and previously unknown individuals not included in the training set. We demonstrate that feature extractors pretrained on species classification can be successfully adapted for this task. Extending individual-level recognition to unknown individuals, so-called out-of-distribution classification, is a crucial step towards making individual recognition a realistic possibility in the wild.

**Highlights:**

- We show that features learned by models pretrained on bird species data can be transferred to individual classification tasks with minimal effort.
- We define and explore the out-of-distribution classification problem on individual animal vocalisations and address various subtleties and eco-logical use cases.
- We compile and contribute two additional datasets to facilitate further research on individual acoustic recognition.

## 1. Introduction

Monitoring individual animals within wildlife populations can facilitate conservation efforts, ecological research and wildlife management. Traditionally, the tracking of individuals has generally been undertaken by physical marking techniques like bird banding [1] or appendage clipping [2]. These individual-level data can then feed into analytical frameworks like mark-recapture studies to estimate population parameters such as population size, abundance, survival rates and movement patterns. However, traditional physical marking approaches have a number of limitations - They are often labor-intensive and costly to implement and potentially harmful to the animals being studied [3, 4, 5, 6]. Furthermore, many physical marking techniques do not endure when the individuals are especially long-lived [7]. In recent years, non-invasive identification techniques have emerged as promising alternatives that address many of the concerns associated with physical marking while still allowing individual-level tracking. This includes photo identification relying on stable external marks or patterns [8] as well as molecular screening of eDNA (e.g. scats or other sources [9]). Bioacoustic monitoring has emerged as an another promising technique and instead uses audio recording devices to capture and analyse animal sounds. The rise of machine learning and deep learning has greatly benefitted the processing and analyzing of the substantial amounts of data generated and deep neural networks are widely applied to automate the acoustic detection and recognition tasks [10, 11, 12].

To date, the majority of work using such audio data has focused on species-level monitoring. This has proven valuable for biodiversity assessments, habitat surveys and tracking broad population trends [8, 10]. Yet, few studies exist that analyse and apply bioacoustic recordings for individual-level tracking [10].

Individuals from numerous species are known to produce individually unique vocalisations that serve a variety of biological purposes including mate recognition, territorial neighbor recognition and parent-offspring recognition [13]. Furthermore, vocalisation characteristics may also differ on the individual-level without any adaptive function, for example, due to anatomy.

In bioacoustic monitoring, these vocalisations have the potential to serve as acoustic fingerprints and may offer a new dimension of insights and benefits for wildlife monitoring [10]. The ability to non-invasively obtain large amounts of individual-level data has the potential to facilitate more accurate population estimates, deeper insights into social interactions and movement patterns of specific individuals without the need for physical capture or tagging.

While there are many promising potential applications, individual acoustic recognition has not gained as much attention as species-level bioacoustic monitoring and remains in its early stages [10, 14]. It requires sophisticated techniques to analyse subtle variations in acoustic features that are often not even noticeable for humans, for example deep learning. Importantly, these methods often require large datasets of labelled calls from known individuals for training, which can be impractical, time-consuming and labor-intensive to obtain. This is especially challenging for wild animal populations, where individual identities are typically not already established. As a result, testing individual acoustic recognition has largely been confined to controlled settings where each individual is known, significantly simplifying the recognition task [10, 14]. These proof-of-concept studies typically involve recording a small, fixed set of animals (e.g. a captive population or tagged subset of the total population) under optimal conditions to confirm that the individual-specific acoustic features can be reliably detected and classified.

The transfer of these methods to natural environments presents significant challenges. Wildlife populations are typically both open and dynamic, with individuals routinely entering and leaving the monitored area. This scenario is significantly more challenging than the closed-world lab setting since, instead of identifying only known individuals, a functioning system must now detect and distinguish vocalisations from both known and unknown individuals. In machine learning this is termed out-of-distribution (OOD) classification. This study explores the challenges of out-of-distribution classification on the individual level. First, we collect and collate two novel datasets for individual acoustic recognition and train a deep learning model to empirically confirm that the vocalisations of these two species are indeed individually distinct and distinguishable by an automatic classifier. Next, we simulate a fluctuating population by withholding a subset of individuals during training and only introducing them during testing. We show that our method can be adjusted to perform OOD classification and to detect these as novel individuals without having been trained on them. Finally, we explore performance improvements and discuss the ecological applicability of our results.

## 2. Related works

Prior research has confirmed individually unique vocalisations in a range of species, primarily birds and mammals [13, 15]. A limited number of studies on automatically classifying individuals has emerged in recent years using a range of different methods including random forests [16], various forms of unsupervised clustering [17], support vector machines [18], neural networks pretrained on images [19] and task-specific neural networks [20, 21]. Notably, the vast majority of studies have focused on a closed-set/in-distribution (ID) scenario (93.3% as surveyed in [10]). The remaining works that focused on an OOD scenario (6.7%) have primarily been performed either through manual classification or using traditional machine learning methods [10], with only a single example leveraging the advances in deep learning [22].

OOD classification has garnered significant attention in the deep learning community due to its ability to handle scenarios where models encounter previously unseen classes during inference [23, 24, 25]. The existing OOD work in other domains presents a compelling opportunity to extend and adapt these methods to individual wildlife recognition applications. Approaches to OOD classification can be grouped into four broad groups [25]: *Classification-based methods* are methods that directly use a trained model’s output, e.g. the class probability to classify OOD samples. The training paradigms for such models can be categorised into those that incorporate a dedicated OOD-specific training objective [26, 27] and those that do not [28, 29]. The former explicitly optimises the model’s performance on OOD detection tasks, while the latter relies on the model’s inherent generalisation capabilities acquired through standard training procedures. Both approaches may incorporate explicit examples of OOD samples during training, disjoint to those used for testing, to facilitate the learning of the ID/OOD separation [30, 31]. Previous work has shown that the efficacy is significantly impacted by the correlations between the OOD samples provided during training and those encountered during testing [32].

*Density-based methods* use probabilistic models to first characterise the ID data and subsequently identify test samples in low-density regions of the modeled distribution as OOD instances. While theoretically sound, these methods often present significant challenges in terms of training and optimisation complexity. As a result, their empirical performance frequently has been found to frequently not surpass that of classification-based approaches [25].

*Distance-based methods* classify samples as OOD *explicitly* based on a distance measure to stored ID samples.^1^ Such methods include those memory-efficient ones that store the centroid or prototypes of ID samples [33] as well as nearest neighbor classifiers, which store all ID samples in memory [34]. The effectiveness of these methods heavily depends on the quality of feature space representation. Many distance-based approaches assume that known classes form tight clusters in the feature space, which may not always be the case. Distance-based methods are impacted by high-dimensional feature spaces, where the difference between minimum and maximum distances becomes indiscernible [35] and irrelevant attributes in the feature space can impede the separability of different distributions. *Reconstruction-based methods* use an encoder-decoder framework trained on ID data and exploit the fact that reconstruction models trained only on ID data cannot recover OOD data well and leverage this discrepancy in model performance to detect OOD samples. In practice, such models are difficult to train due to the problematic trade-off between reconstruction quality and OOD detection capability, where improving one aspect often comes at the cost of degrading the other. This type of architecture faces inherent tensions between compression and feature preservation, while also struggling with scalability issues in high-dimensional spaces and datasets with many categories [36].

One application area of deep learning that already considers OOD scenarios and techniques in an ecological context is the visual re-identification of individual animals [37, 38, 39]. It involves using computer vision and machine learning algorithms to recognise specific animals across different images or video frames, even when the visual conditions or contexts vary significantly from the original training data. The general approach here is to train a feature embedder on ID samples and to compare the test sample embeddings to those of ID samples. Distance-based methods in the form of similarity scores and learned thresholds are then used to either classify the individuals or label them as new, unseen individuals [40, 41, 42].

## 3. Material and methods

### 3.1. Data collection

We used five datasets gathered from open and unconstrained wildlife populations for the performance evaluations of our computational methods in this work (Table 1). The south-eastern red-tailed black cockatoo (*Calyp-torhynchus banksii graptogyne*) and little penguin (*Eudyptula minor*) data are novel and were collected by the authors for this work. The little owl (*Athene noctua*), tree pipit (*Anthus trivialis*) and chiffchaff (*Phylloscopus collybita*) data were obtained from [43].

**Table 1:** Summary of audio data used. While individuals are all known during the closed-set experiments, individuals were split into known and unknown subsets for the out-of-distribution experiments. The unknown individuals were chosen such that their vocalisations total roughly 20% of the total data.

|  | Number of individuals<br>(known/unknown) | Number of vocalisations<br>(known/unknown) |
| --- | --- | --- |
| Red-tailed black cockatoo<br>(female begging call) | 16<br>(12/4) | 1785<br>(1433/352) |
| Little penguin<br>(exhalant display call) | 15<br>(10/5) | 2429<br>(1958/471) |
| Little owl<br>(territorial call) | 16<br>(12/4) | 952<br>(749/203) |
| Tree pipit<br>(song) | 10<br>(7/3) | 1364<br>(1062/302) |
| Chiffchaff<br>(song) | 13<br>(9/4) | 6238<br>(5075/1163) |

The datasets used vary in vocal complexity. Generally, we focus for each species on the call type that is most suitable for individual-level recognition. The red-tailed black cockatoo (RTBC) adult female begging call and the little owl (LO) territorial call consist of a single syllable, while the little penguin (LP) display call consists of an exhalant and inhalant phase. The tree pipit (TP) and chiffchaff (CC) songs are more complex and contain (on average) 11 and 9-24 syllable types repeated in phrases and variably combined respectively [16]. Example spectrograms of the different vocalisations are shown in Figure 1.

**Figure 1:**
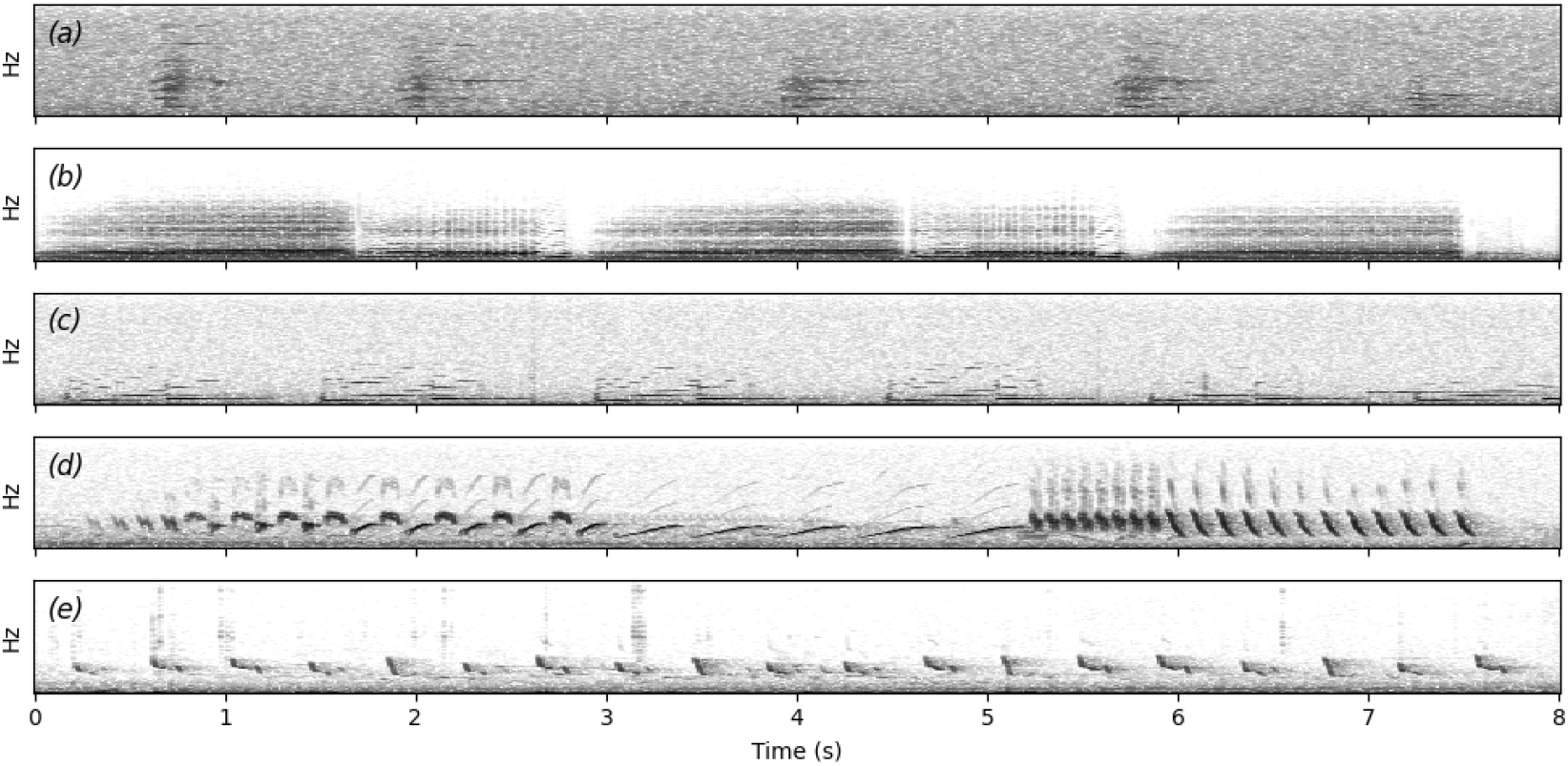
Example spectrograms representing a a) south-eastern red-tailed black cockatoo adult female begging call, b) little penguin display call (3 exhale and 2 inhale components), c) little owl territorial call, d) tree pipit song and e) chiffchaff song.

#### 3.1.1. Little penguins

Little penguins were recorded at St Kilda Breakwater, VIC, Australia (37.86° S, 144.96° E). A combination of Wildlife Acoustic Songmeters Mini and Micro (44.1 kHz, 16-bit, PCM with 6 dB gain) were placed in occupied nests and used to continuously record overnight from 19:00 to 07:00 (all times reported as local time) in the nesting period between April and May 2024. For each nest, we restrict ourselves to audio from a single night (3 nights total) to minimise the chance of recording unwanted individuals (e.g. if individuals moved during the nest site selection phase). The nests between nights were chosen such that they occupied different sections of the breakwater, thus minimising the chance of recording and labelling the same individuals across different nights as different ones. Individual LPs were identified by correlating the vocalisation amplitudes in recordings with the locations of known nesting sites. Calls less than 80% of the maximum amplitude were regarded as originating from neighboring nests or penguins passing by and deemed not clearly identifiable. We note that because the LP nest in pairs and are visually indistinguishable, the vocalisation label we infer is actually a nest-level identity (i.e. a vocalisation from partner 1 or 2 inside nest A is labelled as identity A). However, the distinguishing features of a nest are inherently a combination of characteristics from both individuals and successful nest-level classification would indicate that the combination of traits from individuals in each nest is unique. Consequently, we argue that classification on a nest-level is comparable to classification on an individual-level in the context of determining the ability to classify individuals.

The vocalisations were extracted individually from the continuous recordings and validated by author LH using Raven Pro 1.6.5. We target the exhale component of the display call (see Figure 1b), as it has exhibited signs for individual distinctiveness [44].

#### 3.1.2. Red-tailed black cockatoos

Audio recordings for the south-eastern red-tailed black-cockatoo were obtained for 16 wild nests in South Australia and Victoria. Most of these recordings were acquired from a large, ongoing nest monitoring program run by the South-eastern Red-tailed Black-Cockatoo Recovery Program [45]. Data from two nests were acquired from an existing database of recordings [46]. Recordings used in the current study were collected between 2017 - 2022.

Of the 16 nests, three were natural nests in tree hollows and 13 were nests in artificial nest boxes. Audio recordings were collected using either Frontier Labs BAR recorders (n = 2 nests), AudioMoth recorders (n = 7 nests) or SWIFT recorders (n = 7 nests). Sound recorders were attached to nest trees or, if that was not possible, a nearby structure. Recorders collected continuous audio for 3 – 4 hours each day, commencing roughly 2.5 hours before sunset, which is the time of peak calling activity at nests. Sampling rate varied by recorder type. BAR recordings were made at 44.1 kHz while AudioMoth and SWIFT recordings were made at 32 kHz. All recordings were made in .wav format.

To build the acoustic dataset, sample calls were labelled and extracted from the audio recordings using Raven Pro 1.6 software. Some audio recordings were first processed using a custom call recogniser to detect the species, while others were processed without the aid of any automation. In both cases, audio recordings were manually inspected in Raven Pro by one observer (DT) and vocalisations were labelled to the level of the call type. Only female begging calls were extracted for the current study. Although red-tailed black-cockatoos give many call types at nests [46], the female begging call was chosen as the best proof-of-concept call type because it is reliably given at active nests, which allows sufficient sample sizes of calls, and is not given by males, which reduces the chance of the observer assigning a call to the wrong individual. Begging calls are highly variable within and between individuals, which may or may not aid individual classification. Red-tailed black-cockatoos nest in loose colonies, meaning that begging calls from non-target females may be captured in recordings. To minimise the chance of mislabelling, calls were only extracted if they were loud and clear, indicative of a bird close to the microphone. Because nest hollows may be used by different individuals during a nesting period (e.g. after one nest fledges or fails, another bird may nest there), we only extracted calls for which we were confident that the nest was being used by the same individual.

#### 3.1.3. Little owls

This dataset comprises little owls recorded from sunset until midnight between March and April of 2013—2014 in two Central European farmland areas. The territorial calls were recorded for up to 3 minutes after a short playback provocation (1 min) from a maximum of 50 meters. For full details, see [43, 47].

#### 3.1.4. Tree pipits

This dataset comprises songs of tree pipit males that were recorded throughout the entire day from mid-April to mid-July between 2011 and 2013 in the Czech Republic. Only spontaneously singing males were recorded. For full details, see [43, 48].

#### 3.1.5. Chiffchaffs

This dataset comprises songs of chiffchaff males recorded between 2008 and 2011 in the Czech Republic. Only spontaneously singing males were recorded from within about 5–15 m distance. Because the syllable repertoires of males are reported to differ almost completely between two years [49, 50], we only consider the “within-year” CC data from [43] for the context of this work. For full details, see [43].

### 3.2. Problem formulation

We first show that the individuals from all five species have individually distinct vocalisations in a closed-set (in-distribution) scenario. Consider a labelled training dataset with *N* samples:

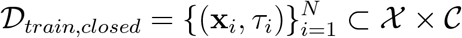

where 𝒳 is the input space and 𝒞 = {1, …*C*} the set of known classes with each class representing one individual. Given a previously unheard vocalisation, the model should return the probability of it originating from a known individual *i*. Formally, given a sample **x** from a test dataset in which all *M* samples are drawn from the same Cartesian product 𝒳 × 𝒞 as the training dataset the model should return a probability distribution over the known classes *p*(*y*|**x**).

Next, we extend our problem to an out-of-distribution scenario by withholding a subset of the individuals during training and treating them as novel individuals during the testing phase. Formally, we split the set of previously known classes 𝒞 into a new set of known 𝒦 and unknown 𝒰 classes:

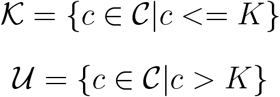

where *K* and *U* = *C* − *K* are the number of known and unknown classes, respectively. The new training dataset is then given by

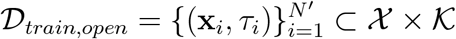

Contrary to the closed-set scenario, the test samples may now also originate from unknown individuals, resulting in

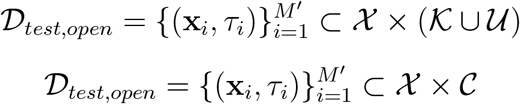

Following the framework defined in [25], we focus on two OOD subtasks with ecological relevance: novelty detection and open-set recognition. Novelty detection requires a model to produce a score 𝒮 (*y* ∈ 𝒦|**x**) for a sample **x** to belong to the known classes 𝒦. A score threshold is then used to classify samples as ODD, i.e. as belonging to either 𝒦 or 𝒰. Open-set recognition additionally requires the model to return a probability distribution over the known classes *p*(*y*|**x**) for all samples assumed to belong to 𝒦.

### 3.3. Closed-set classification

Convolutional neural networks (CNN) have demonstrated robust performance in diverse audio classification tasks, exhibiting their efficacy as a versatile framework for acoustic signal analysis [51]. We train a recent general end-to-end model based on CNNs (AemNet [52]) as the baseline for our closed-set classification experiments. This model skips the conventional spectrogram extraction step and instead converts the raw audio input into a 2D tensor through a series of trainable 1D convolutions. The result is then fed to a classification CNN (Figure 2).

**Figure 2:**
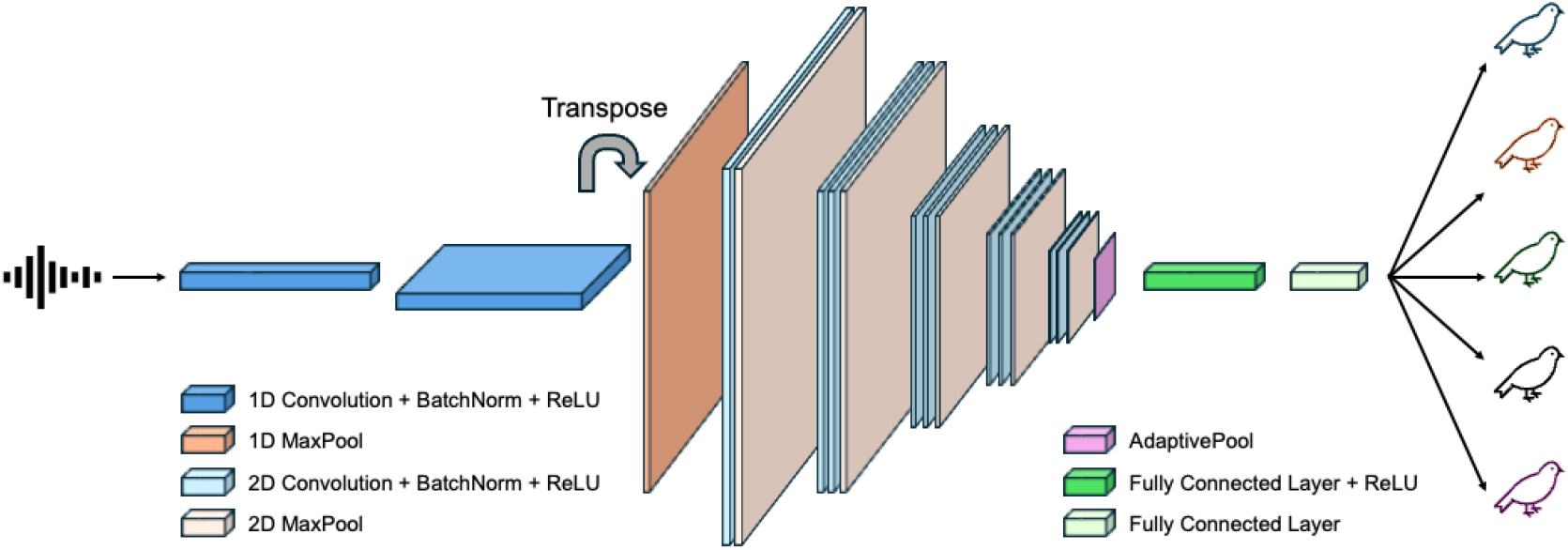
AemNet architecture. The raw audio input is converted through a series of 1D convolutions, batch normalisations, ReLUs and max pooling operations into a 2D tensor, which is in turn fed into a classification CNN.

An adaptive pool in front of the fully connected layers produces a vector of fixed size, allowing the model to incorporate arbitrary length inputs with no architectural changes (the exact architecture can be found in Appendix A). We apply time-masking with a minimum and maximum silent length of 0 and 30% of the total sound length and random gain with a minimum and maximum gain of ± 6 dB. We start the mixup [53] augmentation process after 5 epochs to reduce confusion in the early stages of learning and randomly sample our mixup lambdas from a Beta-distribution with *α* = 5 and *β* = 2. In every epoch, each of these three augmentations is independently applied to a sample with 50% probability.

We then compare the AemNet performance against BirdNET (V2.4) [11] and Google-Perch (V1.4) [12], two CNN-based models widely used for species classification [12, 54, 55]. Although these two models are pretrained on bird species data, they have been shown to perform well on previously unseen bioacoustic classes such as marine mammals and frogs [56]. We use these pretrained models as feature embedders by removing their classification heads and train our own linear classifier^2^ on top of the extracted features.

Because each focal species has varying vocal complexities and call durations, we set the respective input audio lengths for AemNet to be longer than the majority of the individual calls, up to a maximum of 5 seconds: RTBC (2 s), LP (3 s), LO (1 s), TP (5 s), CC (5 s). BirdNET and Google-Perch use constant input lengths of 3 and 5 seconds respectively. Vocalisations that are shorter are randomly zero-padded and longer bouts are cut by selecting a random starting point from which the appropriate length audio is extracted for all models.

All experiments use a 80/20 train/test data split. Note that for the LO, TP and CC data, we deviate from the original work’s train/test split to provide a more consistent comparison between all species results. We use a base learning rate of 0.001, cosine annealing and the AdamW optimiser [57] for all experiments. We train AemNet and the linear classifiers for 200 and 100 epochs respectively. We use five-fold stratified cross-validation for all model training and testing and report the respective average performances. Wilcoxon signed-rank tests are conducted for all relevant experimental comparisons. Given the small sample size of five cross-validation runs, we consider paired samples with a *p*-value of 0.0625 (i.e. the smallest possible *p*-value) as significant.

### 3.4. Out-of-distribution classification

We investigate the OOD classification task on individual animals, focussing on novelty detection and open-set recognition. As with the closed-set classification, we use an 80/20 train/test split, but, as mentioned in Section 3.2, additionally exclude a selection of individuals and their vocalisations totalling approximately 20% of the total data to use as the unknown class (see Table 1). This ensures that at test time, we have approximately a 50/50 split between known and unknown test individuals. We retrain each of the three models on the remaining data using the procedure from Section 3.3 and evaluate their OOD classification performances.

For the novelty detection task, we restrict our experiments to post-hoc methods, i.e. output-based methods that do not need modifying of the training procedure and objective (here: categorical cross-entropy (CE) loss) [25]. This property is particularly important in environments where retraining is either not possible [11] or prohibitively expensive.

We detect novel individuals using two methods: The MSP is a commonly used baseline that uses the maximum softmax probability of a model trained on a closed-set classification as the novelty detection score 𝒮 (*y* ∈ 𝒦|**x**) = *max*_*y*∈𝒦_*p*(*y*|**x**). It has produced state-of-the-art results on various novelty detection benchmarks by leveraging the strong correlation between a model’s closed- and open-set performance [58]. By contrast, the Simplified Hopfield Energy (SHE) method instead uses a simplified version of the Modern Hopfield Network’s energy function to measure the discrepancy between ID and OOD samples [29]. Explicitly, it calculates and stores the average of the trained model’s ID samples’ patterns as 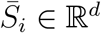:

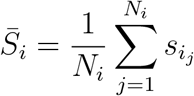

where 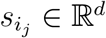 and *N*_*i*_ denote the stored ID patterns and cardinality of class *i* respectively. The similarity of a test sample **x**’s pattern *ξ* ∈ ℝ^*d*^ to the stored averages is then computed using the dot product and used to determine its novelty.

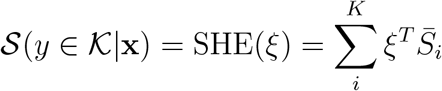

Effectively, SHE is simply the average cosine distance between the sample’s feature vector and all stored feature vectors of the respective class. We use the model’s logits, i.e. the output of the last layer before classification, as the pattern. With its lack of hyperparameters and computational efficiency, this method has resulted in state-of-the-art results on multiple OOD vision recognition benchmarks [29].

Conceptually, it is important to provide the model training with *some* information on what samples outside the known distribution look like if OOD recognition is required. To do so, we adopt Entropic Open-Set loss (EOS) [30, 59]. This loss incorporates additional samples from unknown classes during training to improve the generalisation beyond the closed-set classification task. Intuitively, when presented with an input from an unknown class, a sensible approach is to maintain maximum uncertainty regarding its class membership. This is achieved by enforcing a uniform probability distribution across all known classes, thereby maximising the entropy of the output. We follow the approach in [59] and use a one-hot encoded target vectors *t*_*i*_ ∈ ℝ^*K*^ for each known class *c* ≤ *K*

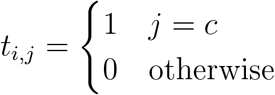

and target vectors with identical values for the unknown classes *i > K*:

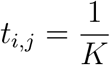

To ensure a diverse coverage of distinctly different sounds, we use the ESC-10 [60] dataset as the negative sample class.

We calculate the area under the ROC curve (AU-ROC) as well as the Open-Set Classification Rate (OSCR) [30, 59] to quantify the novelty detection and open-set recognition performances, respectively.^3^ We require two different metrics, as the AU-ROC considers only the binary classification task and does not provide information on the known class classification performance. The OSCR is defined as the area under the “Correct Classification Rate (CCR) vs False Positive Rate (FPR)” curve, which, similar to the ROC, can be plotted by varying the *θ* threshhold from 0 to 1. The CCR and FPR are given by:

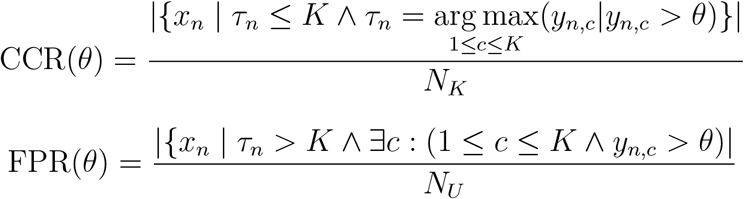

where *x*_*n*_ denotes sample *n, τ*_*n*_ its true class, *y*_*n,c*_ the softmax output probability of class *c* for sample *n*, and *N*_*K*_ and *N*_*U*_ the total numbers of known and unknown test samples. The CCR is the fraction of known samples that have been correctly classified with a softmax probability above a threshold *θ* and the FPR is the fraction of unknown samples that were wrongly classified as a known sample with a softmax probability above the same threshold *θ*. We use five-fold stratified cross-validation for model training and testing and report the respective average performances as well as their standard deviations. Wilcoxon signed-rank tests are conducted for all relevant experimental comparisons. Given the small sample size of five cross-validation runs, we consider paired samples with a *p*-value of 0.0625 (i.e. the smallest possible *p*-value) as significant. Effectively, this *p*-value means that the performance of the better method was superior for *all* cases of the cross-validation.

## 4. Results

### 4.1. Closed-set classification

All three models were able to perform closed-set classification with very high accuracy for all five species (Table 2) on the individual level (RTBC, LO, TP, CC) and on the nest level (LP), respectively. No significant differences were found between BirdNET and Google-Perch on the simple, one-syllable songs (RTBC, LP, LO). The models pretrained on bird species data performed significantly better than the custom model on LP and LO and insignificantly better on the RTBC vocalisations. For the longer TP and CC songs, the custom model and BirdNET performed similarly and significantly better than Google-Perch in both cases.

**Table 2:** Closed-set classification results. For each species, AemNet was trained from scratch, while BirdNET and Google-Perch were finetuned using a two-layer linear classifier. Significant results are shown in bold. Significance is only marked for testing between the respective tested method and the lowest performing method. For example, for Little owls, Perch and BirdNet both perform significantly better than AemNet but no assertion is made about their performance relative to each other.

|  | Model | ACC | AU-ROC |
| --- | --- | --- | --- |
| Red-tailed black cockatoo | AemNet | 95.41±1.10 | 99.91±0.04 |
|  | BirdNET | 97.48±0.18 | 99.95±0.02 |
|  | Google-Perch | 97.25±0.48 | 99.95±0.01 |
| Little penguin | AemNet | 93.63±0.78 | 99.76±0.04 |
|  | BirdNET | **97.03±0.61** | **99.88±0.05** |
|  | Google-Perch | **95.61±0.86** | **99.82±0.05** |
| Little owl | AemNet | 92.04±1.57 | 99.61±0.27 |
|  | BirdNET | **97.59±0.71** | **99.96±0.02** |
|  | Google-Perch | **98.64±0.97** | **99.96±0.05** |
| Tree pipit | AemNet | **94.80±1.45** | **99.80±0.07** |
|  | BirdNET | **92.53±1.82** | **99.55±0.19** |
|  | Google-Perch | 89.30±1.30 | 99.05±0.23 |
| Chiffchaff | AemNet | **97.68±0.21** | **99.96±0.01** |
|  | BirdNET | **97.28±0.48** | **99.96±0.01** |
|  | Google-Perch | 94.92±0.84 | 99.82±0.07 |

### 4.2. Out-of-distribution classification

Our experiments showed that the AU-ROC obtained by using the MSP was consistently higher compared to using the SHE. Furthermore, we found that using EOS loss improved both the AU-ROC and the OSCR across all datasets for both pretrained models. AemNet benefited only in some cases from the inclusion of negative samples and its performance degraded on some datasets (Figure 3). We thus focus on the best combinations of methods for the OOD classification results in Table 3. Complete results are available in Appendix B, Table B.6.

**Table 3:**
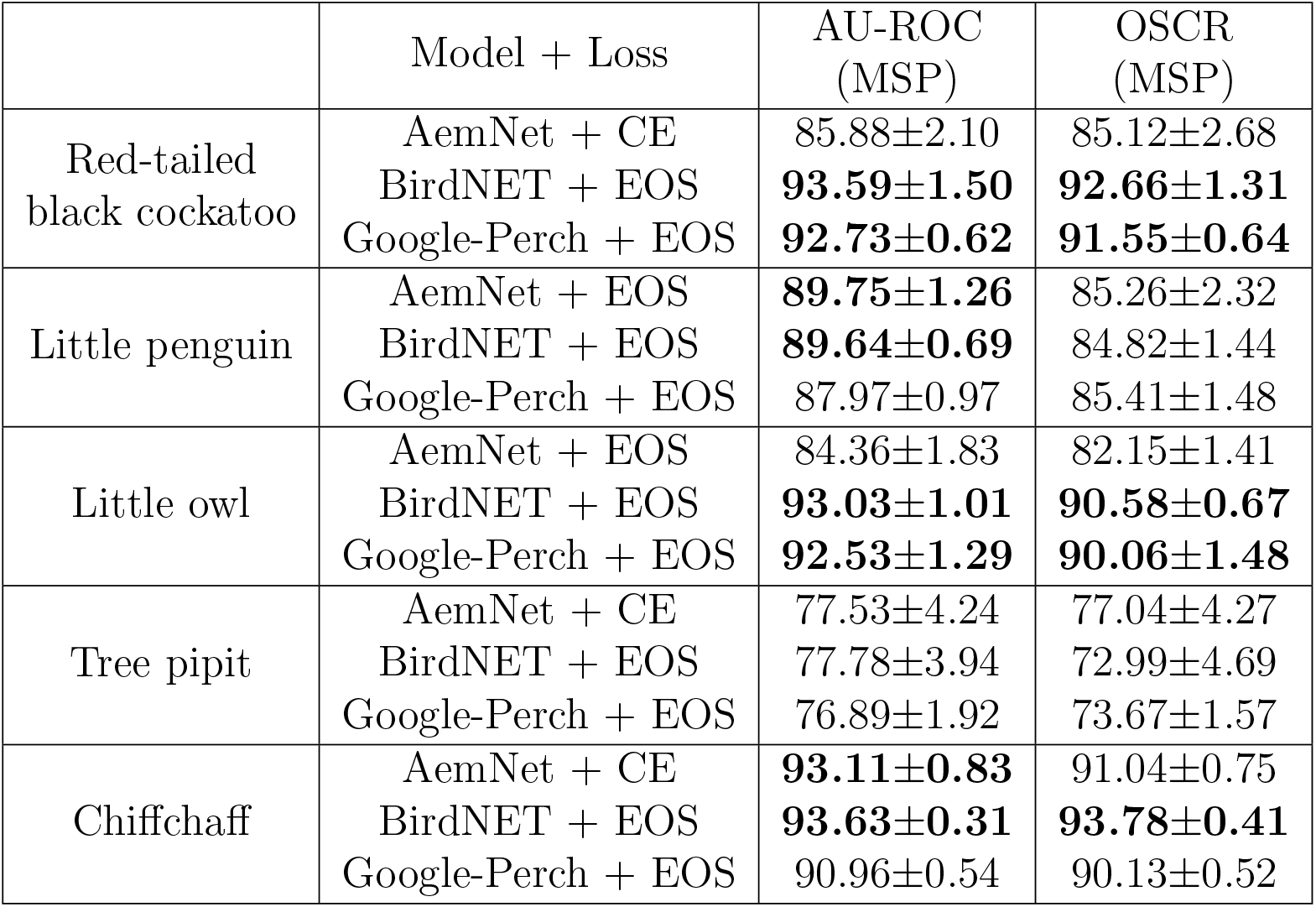
Out-of-distribution classification results. For each species, AemNet was trained from scratch, while BirdNET and Google-Perch were finetuned using a two-layer linear classifier. The best performances for each model and species are reported. Models trained with an EOS loss used samples of the ESC-10 dataset as the negative class. The AU-ROC is reported for the novelty detection task, while the OSCR is reported for the open-set task. Both metrics are reported using the MSP. Significant results are shown in bold.

**Figure 3:**
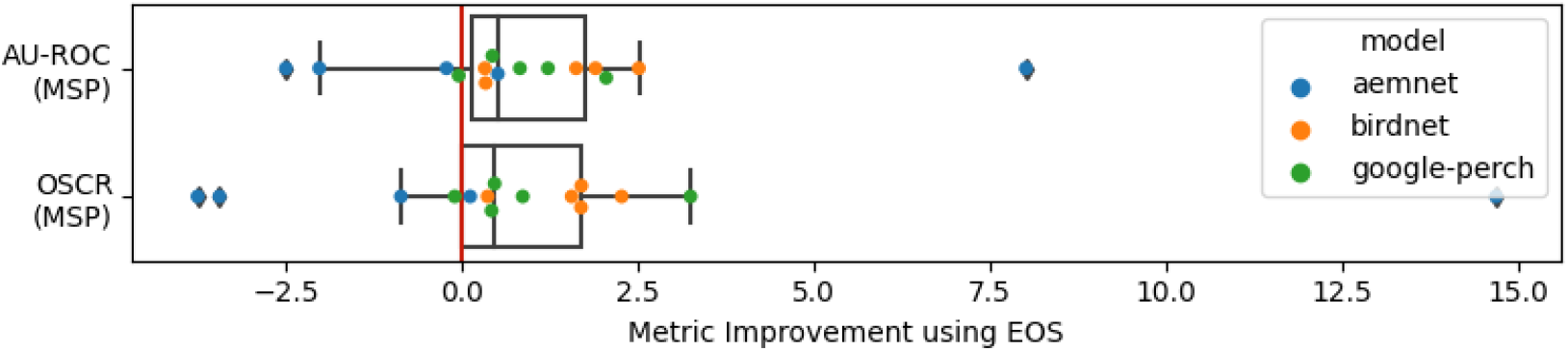
Absolute AU-ROC (MSP) and OSCR (MSP) improvements through training with EOS compared to CE.

BirdNET performed the best across almost all datasets and tasks, while AemNet and Google-Perch were either similar or worse. The more complex TP songs appeared to be the most challenging to classify with lower scores across all models and tasks, with no clear favorite. Considering many of the TP vocalisations were longer than the fixed input length of BirdNET, we repeated our prior experiments with a minor change (Table 4): As we could not increase the input length of BirdNET, we instead decreased the input lengths for both AemNet and Google-Perch to 3 seconds to obtain more comparable OOD classification results (the inputs to Google-Perch were zero-padded to 5 seconds snippets after trimming to ensure model compatibility). Both the AU-ROC as well as the OSCR decreased for Google-Perch and increased for AemNet, albeit insignificantly. Additionally, we retrained our custom model with a 7 second input length (more than 90% of the samples are shorter) and observed a comparable change in performance.

**Table 4:** Pipit out-of-distribution classification results using various input lengths (the 3 second input for Google-Perch was zero-padded to 5 seconds to ensure model compatibiity).

|  | Model + Loss | AU-ROC<br>(MSP) | OSCR<br>(MSP) |
| --- | --- | --- | --- |
| Tree pipit (3 s) | AemNet + CE | 80.33±2.35 | 79.66±2.33 |
|  | BirdNET + EOS | 77.78±3.94 | 72.99±4.69 |
|  | Google-Perch + EOS | 73.30±2.55 | 67.97±1.97 |
| Tree pipit (5 s) | AemNet + CE | 77.53±4.24 | 77.04±4.27 |
|  | Google-Perch + EOS | 76.89±1.92 | 73.67±1.57 |
| Tree pipit (7 s) | AemNet + CE | 79.23±2.84 | 78.80±1.86 |

A positive correlation (Pearson correlation coefficient: 0.78) between the closed-set and the open-set performance was found (Figure 4).

**Figure 4:**
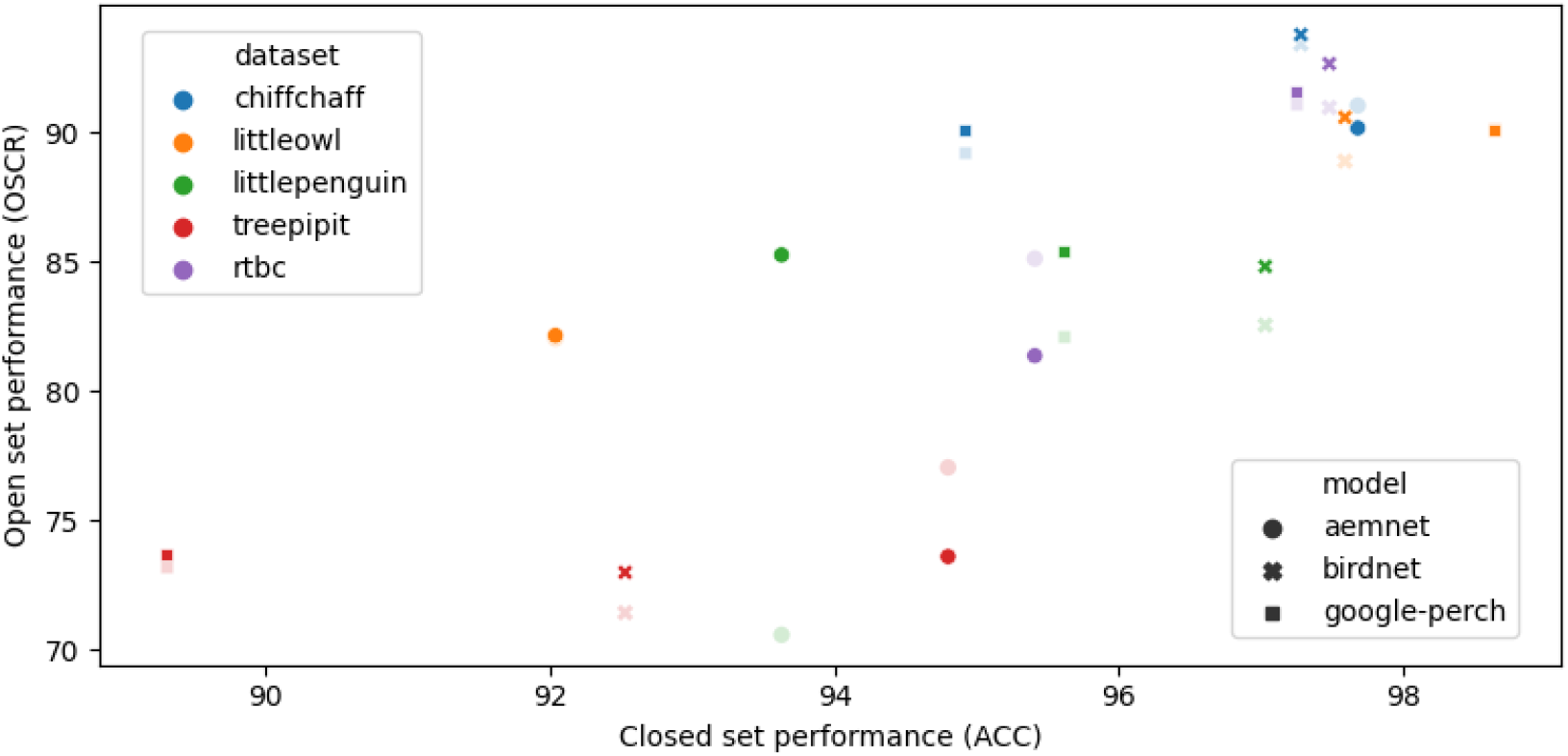
Correlation between closed-set performance (ACC) and open-set performance (OSCR). Results obtained using EOS are shown in bold, while CE results are shown faint.

## 5. Discussion and future work

Our work shows that models pretrained on bird species classification can reliably classify vocalisations of known individual birds and detect unknown individuals across five different bird species. Unknown individuals of species with longer and more complex vocalisations were more difficult to detect, hinting at a limitation of conventional input length models. Nevertheless, our results support the use of pretrained models for individual acoustic recogni-tion tasks.

### 5.1. Closed-set classification

We demonstrated the presence of individually-specific vocal signatures in the red-tailed black cockatoo begging call and exhalant phase of the little penguin display call. This adds to the growing list of taxa with confirmed individually distinct vocalisations and establishes the foundation for subsequent analyses of intra-species vocalisation patterns.

Our experiments reveal an interesting finding regarding the efficacy of pre-trained deep learning models compared to custom-built models for individual acoustic classification. The strong performance of the pretrained models demonstrates that transfer learning from species classification to individual classification works surprisingly well, even without extensive finetuning. This may appear counter-intuitive: During pretraining of these feature embedders, recordings from different individuals of the same species were designated as the same class and differences between individuals thus explicitly neglected.^4^ It is therefore particularly interesting that intra-individual differences were retained in the embeddings generated by these models.

One possible explanation may be that the acoustic features needed to distinguish between species are generally similar to those that vary between individuals within a species. If this is correct, feature embedders trained on a broad variety of species, such as these, can be expected to develop feature sets powerful enough for individual-level classification. An alternative explanation may be that sufficient residual variation between individuals remains in the embeddings per chance, not because this is functionally equivalent to what is needed for species classification. That such variation could be used for classification tasks beyond the species level would be consistent with the fact that even randomly defined features can deliver surprisingly accurate classification results [62].

Notwithstanding these theoretical considerations, the ability to use pre-trained models offers significant advantages from a practical perspective in terms of resource efficiency and accessibility. Not only do pretrained models reduce the computational resources, they are also considerably easier to use and to adapt for non-domain experts without extensive machine learning expertise. This enables a broader range of scientists to leverage deep learning in their acoustic monitoring work, including those working in individual identification.

While the authors in [56] showed that models trained on bird vocalisations learn a wide range of acoustic features generalisable to novel categories, our work demonstrates that these features also generalise to intra-species categories, at least in the case of our test species. As our test species are not distinguished in any obvious way, it stands to reasons that these features may be useful for individual recognition in a much wider range of bird species and possibly other wildlife.

### 5.2. Out-of-distribution classification

Our work differs from the only other work using deep learning in a bioacoustic out-of-distribution setting [22] in several ways. Instead of building and training a task-specific model, we showed that existing models pre-trained on closed-set species classification can be extended to individual out-of-distribution classification tasks with strong performances and minimal effort. We applied this approach approach across more individuals and several species, demonstrating its viability and robustness. We formalized two out-of-distribution classification tasks and used task-relevant metrics to quantify our model performances, allowing for comparison between models as well as improvement suggestions. Our results on these tasks demonstrate two key findings:

First, exposing the models to negative samples during training improved both the AU-ROC as well as the OSCR in the majority of our cases. Interestingly, AemNet’s TP and CC performance declined with the inclusion of negative samples in the training process. This may be due to the fact that AemNet first needed to learn the new feature set of the more complex songs and the negative samples introduced too many confounding factors to an already more difficult task, whereas the pretrained methods were only required to learn a classification task from a known feature set. We note that the classes used were entirely unrelated to the task and that the effectiveness of this approach may be dependent on the nature and diversity of the negative class. As a result, we propose that in addition to targeting labelled individuals during the audio recording process, additional unlabelled individuals should be recorded and included as negative samples during training as well. These are individuals whose exact identity cannot be inferred but are known to be disjoint to the set of known training individuals. This can be achieved e.g. by recording visually different individuals (e.g. unbanded individuals) or recording individuals in a different location (e.g. in a different section of the recording site). Because the EOS loss does not discriminate between the negative classes, the only requirement is that the negative class individuals are not part of the supervised training data. We note that this data is far easier to obtain as no individual labelling is required. This usage of more relevant out-of-distribution inputs may help the models to learn more subtle but relevant differences.

Second, the pretrained models performing comparable to or better than our model trained from scratch on the closed-set task suggests they may have developed more robust and generalisable features for individual identification, which is particularly important for the out-of-distribution classification task. Our results on the simple vocalisations align with this conjecture, however, as in the closed-set case, their performances on the TP vocalisations were similar to or worse than that of AemNet. One reason this may have occurred is that the smaller context window of BirdNET (3 s) may not have been sufficient to cover all of the relevant individuality information of a sample, where the median and average sample lengths are 3.5 and 4.0 seconds respectively. Our results from the extended evaluation of the TP vocalisations (Table 4) showed, however, an upper-bound in performance, suggesting that conventional fixed length architectures may be suboptimal for capturing the full spectro-temporal complexity of long multi-syllable bird songs in the context of individual out-of-distribution classification.

Considering the strong closed-set performances of these models, long-range dependencies important for individuality may have been discarded during training. More flexible computational frameworks that can handle long-range dependencies, such as transformers [63] and structured state space sequence models [64], may be needed to better handle such dependencies within more complex songs.

### 5.3. Ecological applicability

Our results demonstrate robust performance in cases where each individual is known, i.e. closed-set recognition tasks. However, the real-world appli-cability of these findings is limited, as in most cases, conservation monitoring more closely resembles out-of-distribution problems, i.e. scenarios where not all individuals are known.

While the techniques presented in this work provide a solid foundation for identifying known individuals and differentiating between known and unknown individuals, we posit that integrating capabilities to count individuals could substantially enhance the impact of these models. The ability to produce count estimates of unknown individuals in a recording, effectively providing local abundance estimates around a recording device, would be a substantial step forward in estimating population trends from bioacoustic data. This is crucial not only for biodiversity and environmental assessments, but also conservation planning, resource management and decision-making. Source count estimation of unlabelled wild populations presents inherent challenges, primarily due to the number of individuals. The transition from a controlled setting, where the system is exposed to a limited number of individuals, to inference in the wild, where the number of unknown individuals may be much greater, is not straightforward. A system performing well on five unknown individuals in the lab may not be guaranteed to perform well on a large number of unknown individuals in the field. However, addressing these obstacles could yield transformative results.

There are two primary use cases for source count estimation in an out-of-distribution scenario. The first is to confidently identify, or separate, individuals in order to measure abundance at a point in time. Actual identity is not important *per se* and any slow change to vocal signatures over longer time scales is not problematic. The second use case is to identify and label individuals, so that capture-recapture models could be used to measure, for instance, population size. Shift in individual vocal signatures, for example due to ageing and growth, may be problematic in this case. Further study is required, specifically whether domain adaptation techniques can address this.

When individuals are vocally unique, one approach to obtaining source count estimates could leverage the strong performance of pretrained models. As the pretrained feature spaces contain sufficient information to distinguish individuals, it appears likely that clustering in the feature space can be used to (approximately) count individuals. Specifically, deep clustering techniques could be employed, wherein the pretrained embeddings are used to feed into a clustering objective rather than a classification loss. One would expect that similar vocalisations (i.e. those originating from the same individual) cluster together and separate from dissimilar ones (i.e. those originating from different individuals). In an ideal model, the number of clusters generated would represent the number of sources. An alternative to clustering could be density estimation in the feature space. Different to many existing methods of estimating abundance from sound (e.g. cue counting [65]), these methods do not rely on the number of detected vocalisations, but rather the differences in their content. The strength of this approach is that it does not require additional, difficult to obtain (and often unreliable) information, such as cue rates. We intend to pursue this idea in future work by training such a network on our existing labeled dataset and testing its efficacy in the field by comparing this bioacoustic estimate to one obtained by another method, such as cue counting [65], counting using unmanned aerial vehicles [66, 67] or manually counting the individuals within the recording area.

By focusing research efforts on refining and implementing such techniques, we can potentially develop more comprehensive and accurate models that not only classify and separate individuals, but also provide crucial population-level insights. These advancements could significantly enhance our understanding of ecosystem dynamics, support conservation efforts and contribute to more informed ecological decision-making processes.

## Availability

Our data is available at zenodo.org/records/14694741. Our code is available at github.com/lifihuang/individual acoustic recognition ood.

## Acknowledgements

Research on little penguins was conducted under Monash University animal ethics approval SBS 39695 and Parks Victoria access approval AA-0000400. Research on red-tailed black cockatoos was conducted under University of Queensland ethics approval SBS/076/17/VIC.

We would like to thank Flossy Sperring for assisting in the data collection of the little penguin vocalisations. For contributing data for the red-tailed black-cockatoo, we thank BirdLife Australia, the Victorian Government, the University of Queensland and the South-eastern Red-tailed Black-Cockatoo Recovery Team. BM would like to thank Leo op den Brouw for the conversations that inspired this work.

## Author contributions

**Lifi Huang**: Conceptualization; Data curation; Formal analysis; Investigation; Methodology; Project administration; Resources; Software; Validation; Visualization; Writing – original draft **Rohan Clarke**: Conceptualization; Methodology; Project administration; Resources; Supervision; Writing – review and editing **Daniella Teixeira**: Data curation; Writing – original draft **André Chiaradia**: Conceptualization; Methodology; Project administration; Supervision; Writing – review and editing **Bernd Meyer**: Conceptualization; Funding acquisition; Methodology; Project administration; Resources; Supervision; Writing – review and editing

## Appendix A. AemNet architecture

**Table A.5:**
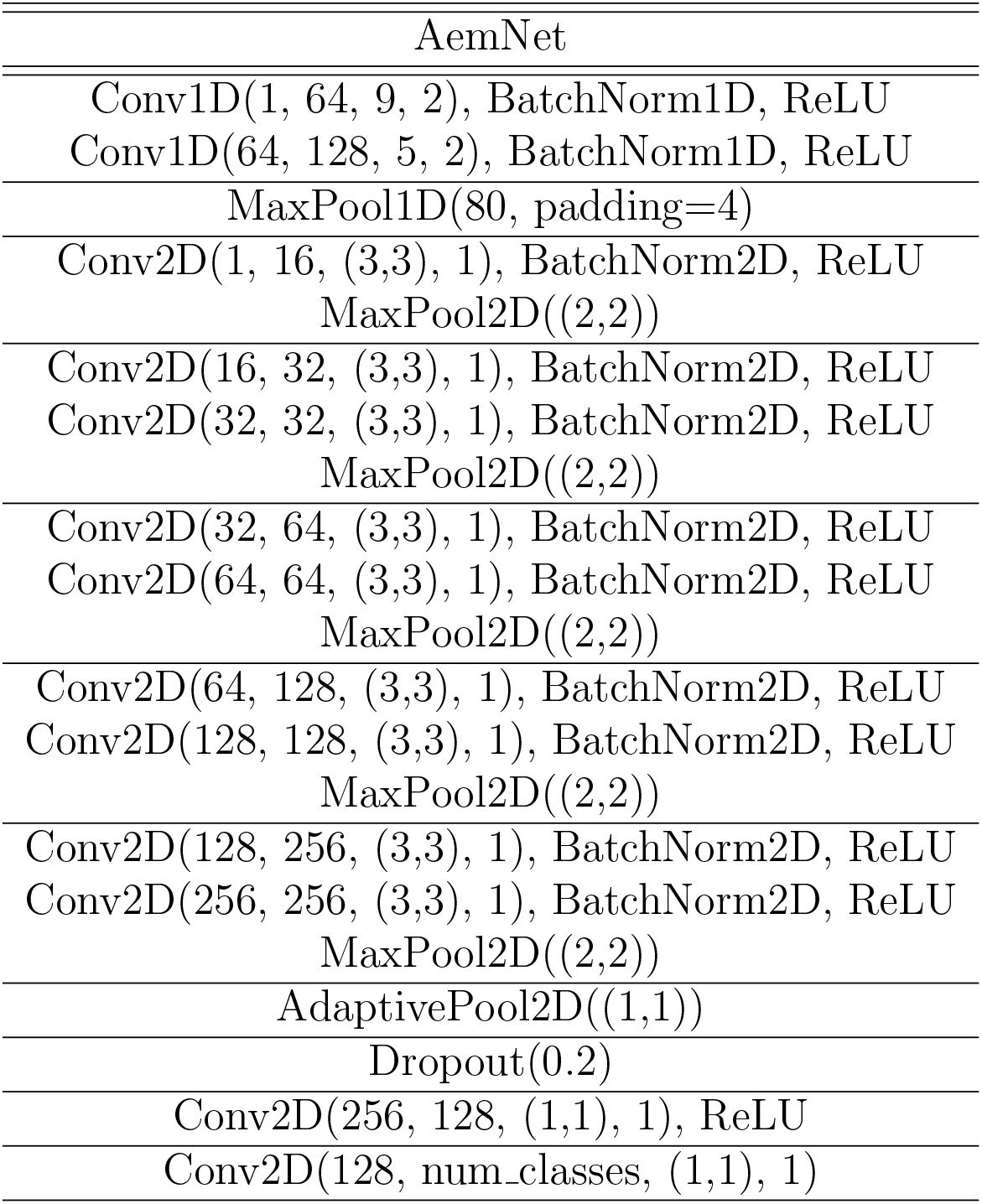
Out-of-distribution classification results. For each species, AemNet was trained from scratch, while BirdNET and Google-Perch were finetuned using a two-layer linear classifier. All models were trained with a CE and an EOS loss using samples of the ESC-10 dataset as the negative class. The AU-ROCs based on the MSP as well as the SHE are reported for the novelty detection task, while the OSCR is reported for the open-set task.

## Appendix B. Further results

**Table B.6:** AemNet architecture. Unless stated otherwise, default PyTorch 2.0.1 argument order and values are used for all functions.

|  | Model + Loss | AU-ROC<br>(MSP) | AU-ROC<br>(SHE) | OSCR<br>(MSP) |
| --- | --- | --- | --- | --- |
| Red-tailed<br>black cockatoo | AemNet + CE | 85.88±2.10 | 63.73±8.78 | 85.12±2.68 |
|  | AemNet + EOS | 83.38±2.85 | 82.62±0.70 | 81.26±2.39 |
|  | BirdNET + CE | 93.27±1.68 | 88.08±6.72 | 90.96±1.48 |
|  | BirdNET + EOS | 93.59±1.50 | 90.89±1.79 | 92.66±1.31 |
|  | Google-Perch + CE | 92.77±0.30 | 91.10±0.95 | 91.13±0.59 |
|  | Google-Perch + EOS | 92.73±0.62 | 90.18±0.55 | 91.55±0.64 |
| Little penguin | AemNet + CE | 81.74±2.47 | 54.31±10.99 | 70.58±3.63 |
|  | AemNet + EOS | 89.75±1.26 | 83.28±4.70 | 85.26±2.32 |
|  | BirdNET + CE | 87.14±0.79 | 55.96±3.27 | 82.55±1.57 |
|  | BirdNET + EOS | 89.64±0.69 | 75.53±2.75 | 84.82±1.44 |
|  | Google-Perch + CE | 85.92±1.07 | 65.03±6.04 | 82.16±0.56 |
|  | Google-Perch + EOS | 87.97±0.97 | 71.12±3.71 | 85.41±1.48 |
| Little owl | AemNet + CE | 83.85±2.83 | 79.59±5.60 | 82.03±3.29 |
|  | AemNet + EOS | 84.36±1.83 | 76.44±4.80 | 82.15±1.41 |
|  | BirdNET + CE | 91.41±1.47 | 87.63±2.43 | 88.88±1.50 |
|  | BirdNET + EOS | 93.03±1.01 | 87.93±3.10 | 90.58±0.67 |
|  | Google-Perch + CE | 91.30±1.81 | 84.11±4.86 | 90.16±1.81 |
|  | Google-Perch + EOS | 92.53±1.29 | 83.35±2.16 | 90.06±1.48 |
| Tree pipit | AemNet + CE | 77.53±4.24 | 77.69±3.79 | 77.04±4.27 |
|  | AemNet + EOS | 75.50±7.17 | 73.37±8.33 | 73.60±6.66 |
|  | BirdNET + CE | 75.89±2.17 | 68.22±1.54 | 71.43±2.96 |
|  | BirdNET + EOS | 77.78±3.94 | 69.41±2.44 | 72.99±4.69 |
|  | Google-Perch + CE | 76.07±2.19 | 75.99±1.26 | 73.21±1.29 |
|  | Google-Perch + EOS | 76.89±1.92 | 76.62±3.33 | 73.67±1.57 |
| Chiffchaff | AemNet + CE | 93.11±0.83 | 91.40±2.74 | 91.04±0.75 |
|  | AemNet + EOS | 92.89±1.01 | 90.64±1.99 | 90.17±1.47 |
|  | BirdNET + CE | 93.29±0.31 | 91.68±0.72 | 93.41±0.22 |
|  | BirdNET + EOS | 93.63±0.31 | 91.32±1.00 | 93.78±0.41 |
|  | Google-Perch + CE | 90.53±0.50 | 91.51±0.51 | 89.26±0.48 |
|  | Google-Perch + EOS | 90.96±0.54 | 91.21±0.88 | 90.13±0.52 |

## Footnotes

1 Note, however, that methods in the other classes may also implicitly involve the use of distance measures.

2 The linear classifier consists of two linear layers separated by a ReLU: Linear(input dim, 256), ReLU, Linear(256, num classes). The input dimension to the first linear layer corresponds the dimension of the feature embeddings (BirdNET: 1024, Google-Perch). Unless stated otherwise, default PyTorch 2.0.1 argument order and values are used.

3 We use the roc auc score function from scikit-learn 1.3.0 to compute the AU-ROC. To compute the OSCR, we first calculate the CCR and FPR for *n* equidistant *θ*-values ranging from the lowest to the highest experimental softmax probabilities, where *n* is the total number of test samples. We then use the trapezoidal rule to compute the area under the CCR/FPR curve, i.e. the OSCR.

4 Both pre-trained models were primarily trained using Xeno-Canto data [61]. While Xeno-Canto contains audio samples for all our five target species, only 16 LP and 74 RTBC recordings were available at the time of pretraining, while over 1000 recordings were available for LO, TP and CC. The Xeno-Canto LP recordings do not distinguish between the inhalant and exhalant phase and the RTBC data labels do not specify the call types, as the majority of recordings were uploaded before the call types were defined in [46].

